# The sea urchin *Paracentrotus lividus* orients to visual stimuli

**DOI:** 10.1101/2024.01.05.574409

**Authors:** John D. Kirwan, Tianshu Li, Jack Ullrich-Lüter, Giancarlo La Camera, Dan-Eric Nilsson, Maria Ina Arnone

**Affiliations:** Stazione Zoologica Anton Dohrn, Naples, Italy; Arquimea Research Centre, San Cristóbal de la Laguna, Spain; Center for Neural Circuit Dynamics, Stony Brook University, NY, USA; Museum für Naturkunde, Berlin, Germany; Lund Vision Group, Dept. of Biology, Lund University, Sweden

**Author notes:** **Summary statement** The behaviour and mathematical modelling of Paracentrotus lividus object taxis indicate that sea urchins typically have coarse spatial vision facilitated by dispersed photoreceptors feeding into their decentralised nervous system.

**Keywords:** Paracentrotus lividus, Vision, Neurosensory Model, Visual Behaviour, Spatial Resolution, Neuroethology

## Abstract

Though lacking eyes, some sea urchins can see: Several species exhibit resolving vision, as distinct from mere light detection. How and where light is captured in the eyeless sea urchins, and how this information is integrated to elicit visual behaviour, remains a fascinating enigma. We assessed the spatial resolution of the sea urchin *Paracentrotus lividus* in laboratory experiments using fifty adults from the Bay of Naples. This keystone species is an important grazer of the NE Atlantic and Mediterranean and a model system to study development.

We carried out behavioural trials in which individuals were placed in a submerged cylindrical arena to determine if they orient towards a visual stimulus on the arena wall, under diffuse, downwelling light. We adopted a novel isoluminant stimulus, necessitating vision of a given resolving power around the horizon to be detected. We tested individuals at five stimulus widths, including a uniform control. Animals oriented (upon clearing an obstacle) only to the widest stimuli (45 deg and above). This acuity may suffice for tasks such as finding nearby shelters or distant patches of habitat.

We modelled the visual and neuronal processes to recapitulate these responses in *P. lividus*, by fine-tuning the model of Li et al. (2022), as applied to the sea urchin *Diadema africanum*. While these species differ morphologically, the model robustly predicts angular sensitivity in keeping with the behavioural experiments. We find that *P. lividus* (and likely many Echinacea) possesses coarse spatial vision and that the neurosensory model applies broadly to sea urchins.

## Introduction

The ability to detect light and visually resolve objects constitutes an evolutionary advantage for many organisms. This is obvious for fast-moving predators and visual specialists with large eyes and centralised brains, such as raptors or octopuses, but is also true of animals with rudimentary vision and simpler visually-guided behaviours. Several species of sea urchin - marine invertebrate grazers lacking discrete eyes or a centralised brain - also possess spatial vision, although with dramatically worse acuity (Kirwan et al. 2018; Yerramilli and Johnsen 2010, Sumner-Rooney & Ullrich-Lüter 2023). Their proposed mode of non-ocular vision entails resolving an image using distributed photoreceptors feeding into the animal’s decentralised nervous system.

Several sea urchin species have been reported to approach dark targets presented along the outer edge of a behavioural arena, which has been interpreted as visual orientation (reviewed by Sumner-Rooney & Ullrich-Lüter 2023). Based on early findings that the body wall of sea urchins is generally photosensitive (Yoshida & Millott, 1959; Millott & Yoshida, 1960) two different mechanisms have been proposed, as to how sea urchins can visually resolve objects. Woodley (1982) proposed that spines shading the oncoming light at varying angles might allow the sea urchin *Diadema* to visually resolve objects.

Behavioural studies expanded on that proposed model by measuring species-specific spine density and spine spacing and correlating those to their findings of visual acuity in several regular sea urchin species (Blevins and Johnsen, 2004; Yerramilli and Johnsen, 2010; Jackson and Johnsen, 2011). However, spine density, spacing and the visual acuity are not generally correlated across sea urchin species (Notar et al. 2022). Spatial expression patterns of visual photopigments have challenged the *spine-shading* model of sea urchin vision. An R-opsin photopigment, which mediates vision in many invertebrates, including the related sea stars (Lowe et al., 2018), clusters distinctly within sea urchin tube feet, rather than being dispersed across the body surface (Ullrich-Lüter et al., 2011, Lesser et al., 2011, Agca et al., 2011).

Once light is captured, the decentralised nervous system must process this sensory input and generate coordinated behavioural output. Li et al. (2023) have recently proposed the first theoretical model for visually-guided behaviour in a sea urchin, based on visual behaviour in *D. africanum* (Kirwan et al., 2018). This model predicts responses to a wide array of visual stimuli in this species.

*Diadema* species exhibit both a clear and robust startle response (spine convulsions in response to a loom), and object taxis (visually-guided locomotion) and have been studied behaviourally and physiologically. Yet, they are phylogenetically distant from most regular sea urchins (Mongiardino Koch et al., 2022) and differ morphologically. *Diadema* has long hollow spines upon a rigid test, and walks by levering itself with large, robust spines. Most regular sea urchins grasp with their mobile, extendable tube feet for locomotion, while *Diadema* has mostly small, unsuckered tube feet (Randall, 1964). This renders *Diadema* an imperfect model to predict behaviour in most regular sea urchins.

We chose *Paracentrotus lividus* (Lamarck, 1816) (Fig. 1) as an experimental model to test their reaction to visual stimuli and simultaneously create a species-specific theoretical model which can generate testable predictions. *P. lividus* is a sea urchin found in the NE Atlantic and Mediterranean, which possesses ambulacral photoreceptors and is typical of regular Euechinoidea in its size, morphology, and tube-foot locomotion.

**Figure 1:**
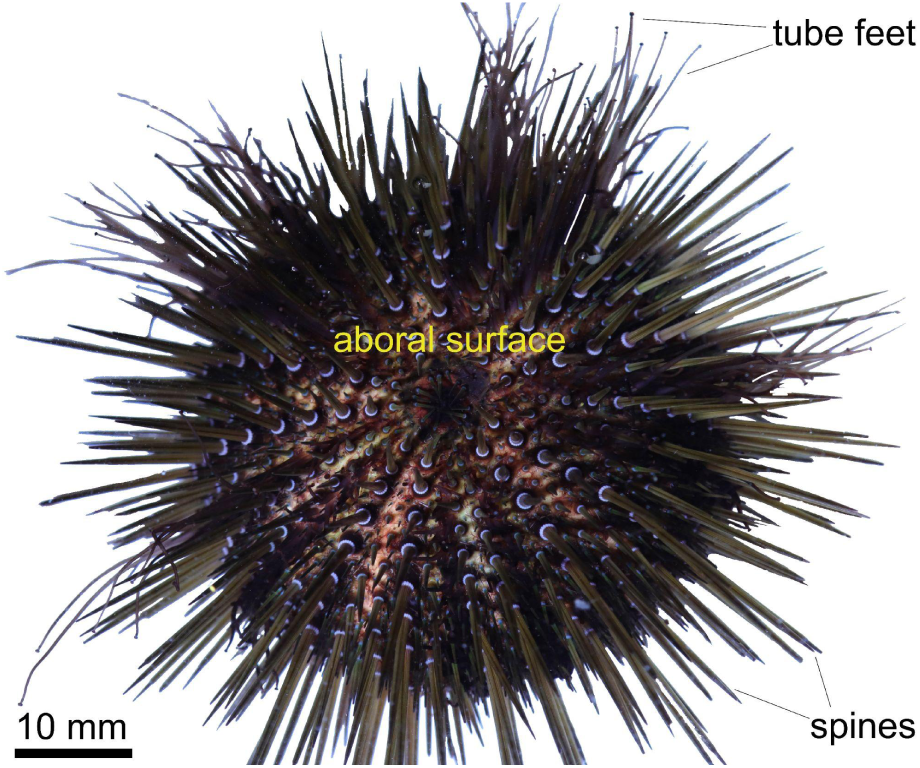
Photograph of *Paracentrotus lividus* adult from above.

We used behavioural arena experiments to test the resolving power of *P. lividus* vision. To clearly discriminate between the animals’ ability to spatially resolve objects versus to detect the reflection of different illumination levels from a stimulus pattern along the arena circumference, we used a 1^st^ Hermitian wavelet as a stimulus. We applied the mechanistic model of sea urchin vision of Li et al. (2023) to the behavioural responses of *P. lividus*.

## Methods

The behavioural experiments measured an object taxis response of *P. lividus* to visual stimuli, to demonstrate spatial vision. The experimental setup resembles that of Kirwan *et al*. but exceeds it in all dimensions, allowing for more stable temperature, water clarity and cleanliness. The angular size constancy of the stimulus is also improved, i.e. for a given distance of travel across the arena floor, the change in apparent size of the stimulus for an observer is lessened.

### Animal husbandry

Fifty adult *P. lividus* were collected from the Bay of Naples. We measured the combined environmental light field of one site, Marechiaro (Fig. S1), as per Nilsson & Smolka (2021). The sea urchins are likely to be three years or older, based on body size and of undetermined sex; all were used in the experiments.

They were kept in a 500 L flow-through plastic tank at Zoologica Stazione Anton Dohrn. Shallow seawater pumped from the bay, filtered and UV treated. They were fed *Ulva sp.* twice weekly and sweetcorn weekly. Within the tank, animals were kept submerged in floating plastic containers, separated into numbered mesh partitions. They were kept on a 12:12 light cycle under fluorescent ceiling lights. The water temperature fluctuated with that of the bay, from approximately 12° to 29° C over the course of the year.

### Behavioural setup

The experimental chamber comprised a further 500 L plastic tank, and adjacent to the animals’ housing. The tank was 93 cm wide, 110 cm long and 70 cm high internally and covered with a wooden lid. Vibrations from outside were dampened by placing ¼″ (6 mm) sorbothane sheets (Thorlabs, Newton, NJ, USA) below the feet. Prior to experiments, the chamber was filled up to 50 cm with natural seawater. Experimental trials were conducted in an arena comprising a cylinder of poster paper, enclosed by opaque black paper atop a wooden base. The circular arena within the tank was 90 cm in diameter.

### Lighting

The arena was illuminated by a pair of lights (Radion XR30W G4; Ecotech Marine, Bethlehem, PA, USA) juxtaposed above the arena to present a square of LED clusters above the centre of the arena, which provided broad spectrum illumination. Below this, a diffusion filter (434 Quarter Grid Cloth or 250 Half White Diffusion; Lee Filters, Andover, UK) covered the arena to mitigate axial light radiating the arena floor unevenly or other visual cues. The diffuser comprised two sheets of 3/4 diffusion paper (Lee Filters 416; 50% transmission at 400–700 nm) between sheets of Makrolon^®^ (Covestro, Leverkusen, Germany) polycarbonate suspended below the lamps on fishing line.

### Stimuli

Previous behavioural experiments (Kirwan *et al*. 2018) with urchin vision applied different visual stimuli. These previous stimuli included a difference of Gaussians (DoG) of opposite amplitude, for which the variance (σ) of the dark Gaussian was half that of the light one. This produced a central dark region with a black maximum, flanked by lighter regions with white minima. The averaged luminance over the complete stimulus equals that of a corresponding subtense of the background, i.e., it is isoluminant.

Here, we have adopted a novel visual stimulus on the arena walls: they were shaded according to a 1^st^ Hermitian wavelet, which is the first derivative of a Gaussian function with μ=0, σ=1:

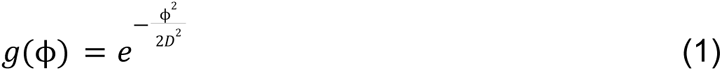

with

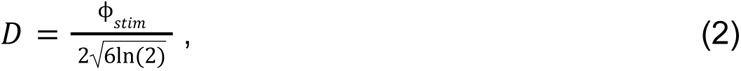

and

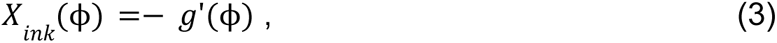

Where Φ_stim_ is the angular width of the stimulus (equal to 0, 15, 30, 45, and 60). These values were calibrated by the curve in Fig. S2 so that the final ink values result in a reflectance inversely proportional to *X_ink_*(Φ).

The 1st Hermitian stimulus comprises a dark and an adjacent light region of equal amplitude (against an intermediate background at both flanks; see Fig. 2C). With perfect sampling, two neighbouring sampling units, whose midpoints are separated by an angle of half the stimulus width and directed towards the stimulus optima would detect a large intensity difference, resulting in perfect Michelson contrast in the case of a totally black minimum. Our printed stimuli had 4% reflectance for the black minimum, resulting in a Michelson contrast of 0.93.

**Figure 2.**
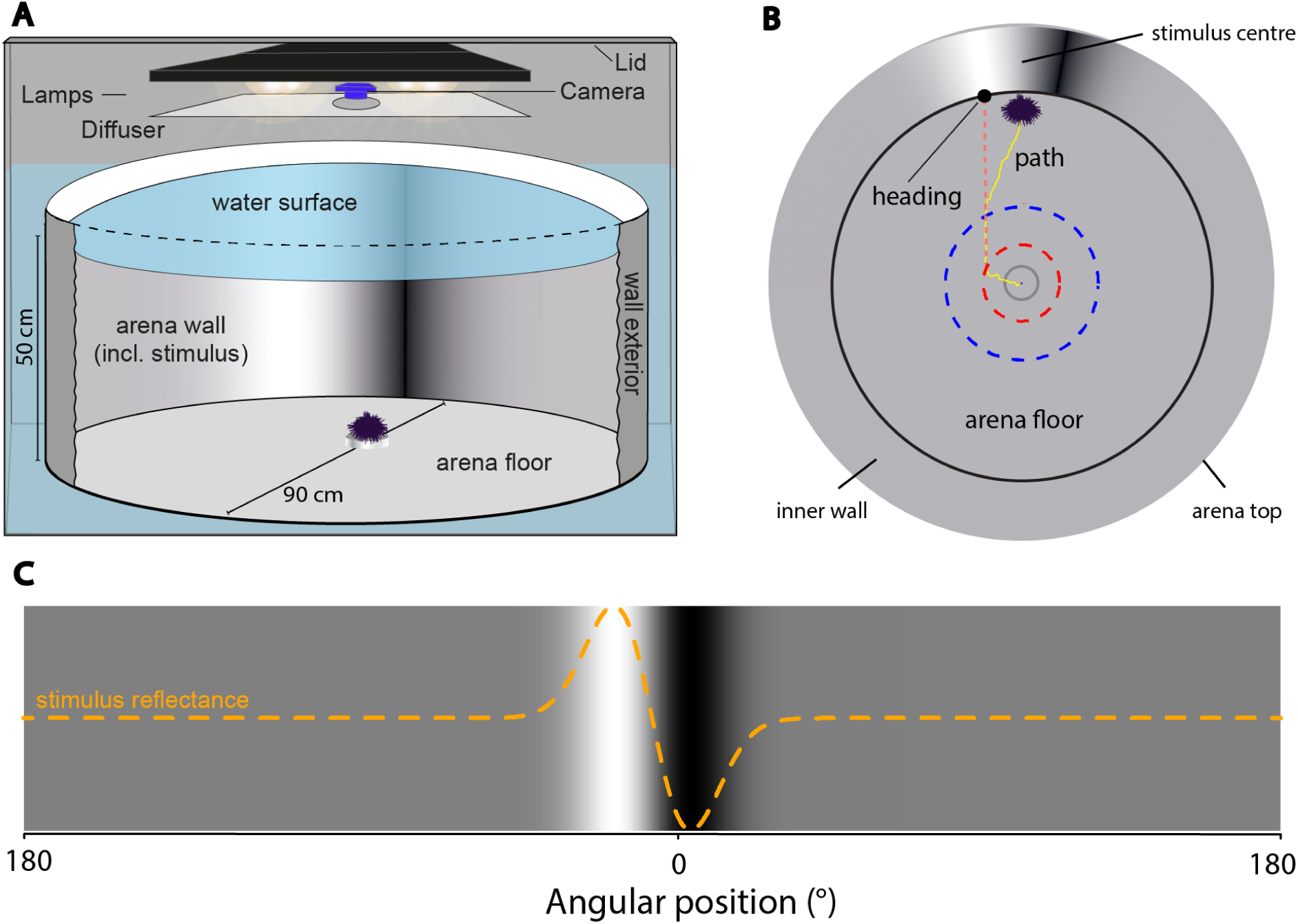
Behavioural setup to assess visual response to a printed stimulus around the horizon in *Paracentrotus lividus.* A. Sectional view of behavioural chamber used in trials. B. Top view of the arena from within the chamber; during trials, the sea urchin moves from the arena centre, its path is tracked relative to the stimulus centre. The dashed line indicates how the heading is found: by the intersection with the arena wall (relative to the stimulus) of a vector of the two tracked points nearest to a fifth and two fifths of the arena radius from the centre. C. The patterned arena wall showing the stimulus in the centre. The relative reflectance across the wall is overlaid as an orange dashed line.

The (angular) resolution of vision is conventionally expressed in cycles per degree which signifies how many periods of the fundamental spatial frequency of the detected waveform occur over one degree of angular width. We express resolution in degrees, as there are far fewer than one cycle per degree. For a sine wave, resolution is the reciprocal of the sole spatial frequency.

The novel wavelet stimulus has several properties which enable it to elicit liminal responses to a shaded object. The 1^st^ Hermitian stimulus contains few higher spatial frequencies (i.e., finer details) above the fundamental frequency. This differs from stimuli with a local high rate of change and especially discrete edges, such as the dual black and white bars of a Haar wavelet. It provides adjacent, anti-symmetrical regions of light and dark relative to the background without discrete edges. Edges comprise further higher spatial frequencies (and therefore greater uncertainty as to which frequencies are detected), albeit these finer superluminal frequencies will go undetected at stimulus widths below threshold.

### Material

Shaded patterns were produced (Coated Fogra39, high quality print PDF, 150 ppi) using Adobe Illustrator (San Jose, CA, USA). They consisted of rich black printed images, which were uniform in the vertical plane but in the horizontal plane had reflectance corresponding to the stimulus, set against an intermediate grey background. The patterns were printed on polyvinyl chloride fabric (EcoFlat 398 gsm blackout banner; Pixartprinting, Venice, IT) using CMYK inks as proportions of rich black (FOGRA39).

To match the shading to reflectance, the side-welling radiance reflected from the patterns was measured within the complete setup with a calibrated TriOS Ramses spectroradiometer (TriOS, Germany), placed 5 cm from the arena wall. Background luminance was below 0.1 lx. The centre of the lens was raised 40 mm from the arena floor and parallel. Relative spectral irradiance (in 3 nm bins) of test pieces of material were measured in the arena setup with the same lighting regime and the wavelength range 425 to 575 nm. The shade of 20 samples of shaded paper of 12.5 cm^2^ was measured at a range of proportions of ink. We normalised these to the highest value to get relative available irradiances for each shade, then modelled changes in available light relative to ink value and printed the patterns accordingly (Fig. S2).

### Obstacle

The animals were placed atop a clear acrylic cylinder, which partially obstructed their locomotion in all directions, but not their field of view. In preliminary testing, we used a wide, salient stimulus, of 150 deg (Fig. S3; 105 trials with some repeats per animal) to check for evidence of a response before testing with a range of narrower stimuli). This evidenced only a small effect, when animals were placed on the arena floor centre, as most animals began a straight and apparently arbitrary course to the wall, shortly after placement. However, we observed that on reaching the arena wall, animals frequently settled close to the stimulus, and we sought ways to motivate them to orient during trials. We found that when the animals are partially obstructed, they were more likely to orient and approach the stimulus.

### Recording

An 3MP USB video camera (Ailipu Technology Co., Shenzhen, China) recorded videos of the trials. A 170 deg lens captured the arena floor and part of the walls. We positioned the camera lens above the water surface, fixed to the LED clusters and directed towards the centre of the arena floor. Videos were shot at 60 fps with a minimum of 720p display resolution and thinned to 5 fps using ffmpeg (https://ffmpeg.org/).

### Tracking

We tracked individual sea urchin paths as per Kirwan et al. (2018) using dtrack (https://bitbucket.org/jochensmolka/dtrack) within MATLAB (MathWorks, Natick, MA, USA). The arena centre and dark maximum of the stimulus were marked in each recording as well as the centre of the animal at each frame. We found the vector of the two tracked points nearest to a fifth and two fifths of the arena radius from the centre, respectively, to represent the path from the centre towards the wall. The heading constituted the angle formed between the intersection of the vector with the arena wall and the stimulus centre. An animal moving in a straight line towards the stimulus centre would have a heading of 0 deg and in the opposite orientation would have a heading of 180 deg.

A trial was completed if the animal moved at least 35% of the radial distance from the arena centre. Occasionally, individuals would remain at or near the centre of the arena, and such trials would need to be repeated (this was especially true of larger individuals). At each trial, the arena wall (which includes the stimulus) was rotated to a position chosen by randomly generating a degree of arc to position the stimulus centre relative to a fixed position on the tank wall.

### Dichotomization

Even when obstructed and with a large stimulus, most animals did not approach the stimulus. This was in keeping with previous trials with sea urchins (Kirwan et al. 2018), but in contrast to a robust visual response observed in a brittle star (Sumner-Rooney et al. 2020). We were therefore minded that the analysis should allow a considerable lapse rate. Behavioural experiments in circular arenas are sometimes evaluated with classical circular statistical tests, which do not account for large lapses. Moreover, while the data is distributed with circular support, we were ultimately interested in whether the animals oriented to the stimulus and not in giving differential weight to directions of travel which are clearly disoriented.

We dichotomised the headings into successes or failures of detection by whether they oriented to a 72 deg sector of the arena wall containing the stimulus, this sector being large enough to contain many headings by chance but not so wide as to dwarf the stimulus, in most cases. We then applied a nonlinear logistic regression model incorporating the psychometric function to these dichotomised data, as per Kirwan and Nilsson (2019).

### Statistical analysis

To find the spatial resolution, we modelled the responses using Bayesian estimation in the Stan language (Stan Development Team, 2023) via the *brms* package (Bürkner, 2017) within R (https://cran.r-project.org/). We used a logistic regression model of the psychometric function with estimated upper and lower asymptotes to represent the chance and lapse rates of the response respectively. The chance rate is the rate at which successful orientation occurs at random in disoriented animals and the lapse rate is that at which animals fail to respond to a maximally detectable stimulus of this type. Fifty independent replicates per stimulus level (i.e. trials with different individuals) was chosen to be enough that, even with a high lapse rate, an effect could be detected.

The criterion used to define the estimate was the inflection point of the psychometric function curve. The psychometric function is a description of the relationship between performance (in this case, visual stimulus detection) and stimulus intensity (the arc width of the difference of Gaussians stimulus) (see Knoblauch & Maloney, 2012; Wichmann & Hill, 2001). This was used because it is a feature of the fitted curve and not dependent on the bounds of an arbitrary interval and is commonly used as a measure of detection threshold (i.e. it is a parameter of the fitted curve that is relatively robust to sample size). The psychometric function *x* was formulated following Houpt and Bittner (2018):

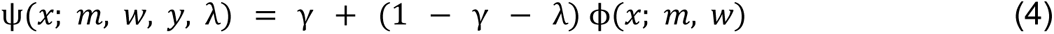

where γ is the chance rate of the response and λ the lapse rate, represented by lower and upper asymptotes, respectively. Φ(*x*; *m*, *w*) is the joint distribution of parameters was modelled according to a sigmoidal logistic function, where *x* is the stimulus width, *m* is the threshold of detection and *w* is the difference between the stimulus width which would produce 10% success versus that which would produce 90% success if γ and λ were 0. The model is defined such that:

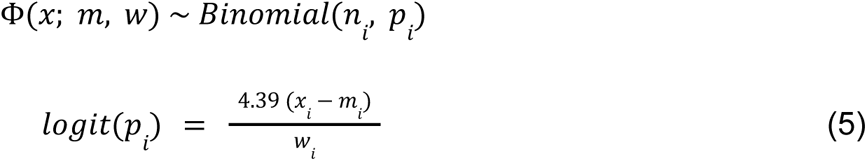

where logit is the logistic transformation, n is sample size, and p is probability. Informative priors (see Fig. S4) were used on λ to facilitate convergence, on x and w to favour a threshold in the range of stimulus sizes applied, on γ to constrain it close to the random probability of success (1/5). The model was run for 4 000 post-warmup iterations from four Markov chains and all posterior draws were analysed.

We depicted the model by examining the influence of the stimulus arc width. The median fit line is plotted (Fig. 4), alongside the 95% credible interval of the draws, which indicate a 95% chance that the true effect lies within this range as per the model. To validate that our model, which suggests a correlation between increasing stimulus size and response, outperforms one without this correlation, we used approximate leave-one-out cross-validation. This method assesses prediction accuracy in unseen data and is suitable for Bayesian models (Vehtari, Gelman, & Gabry, 2017).

We additionally carried out binomial tests of successful orientation for each treatment, using R. This tested the null hypothesis that the proportion of animals directed towards the target sector should be equal or less than would be expected by chance for disoriented animals. For null-hypothesis tests, 0.05 was the critical value (α). We corrected p-values using the false discovery rate (Benjamini & Hochberg, 1995).

### Neurosensory computational model

The behavioural results were captured by a model of decentralised vision in echinoderms recently published in Li et al. (2023). Briefly, the model consists of three components (see Fig. S5): photoreceptor cells (PRC) located on the tube feet, five radial nerves (RN) along the ambulacra, and one oral nerve ring (ONR) surrounding the mouth. For simplicity, neurons with similar properties in each category were grouped together. PRC groups send inhibitory input to their neighbouring RN groups on the same ambulacrum; the latter activate entry stage inhibitory ONR neurons (iONR), which in turn will inhibit output excitatory ONR neurons (eONR). eONR neurons are laterally connected to each other so that their collective activity can spread along more than one ambulacrum (Fig. S5).

Vision in this model depends on the properties of the PRCs and the neuroanatomy of the whole nervous system. The PRC system serves as the input stage since it receives the incoming light; the output of PRCs is further processed by the RNs, whose neural activity is finally integrated in the ONR; the latter serves to produce vision as well as orienting behaviour. The fraction of incoming light received and transmitted by the PRCs depends on the overall acceptance angle of the PRC system. Each PRC had an individual acceptance angle (i.e., the full width at half maximum of their normalised angular sensitivity curve) of 30°, but the PRCs themselves were located uniformly on the tube feet, spanning an overall 30° width along the direction orthogonal to the ambulacrum (see Fig. S5A, where δ is the half-width of the location distribution of the PRCs along the longitudinal angle ϕ). When combining these two features together, an ‘effective’ acceptance angle of 60° results for the overall system (Li et al., 2023).

Upon reaching the ONR, the visual input processed by RN neurons elicits a profile of collective activity in the eONR neurons. The activity profile of the eONR neurons is read out as a ‘population vector’, which is the vector sum of the preferred orientations of each eONR neuron weighted by their own activity level. This way, the population vector points in the direction of the most active eONR neurons, which tends to be highly congruent with the direction of the ‘centre’ of the stimulus (the region around 0° in Fig. 2C). When the length of the population vector exceeds a threshold, the visual stimulus is detected. Full details of the model are in Li et al. (2023).

The stimuli were the same as in the experiments (Fig. 2C), and provided an input current to the PRCs given by *X*_0_(ϕ) = 0. 96(1 −*X_ink_*(ϕ)) + 0. 04, where *X_ink_*(ϕ) was the value used in experiments (but normalised between zero and one). The value of *X*_0_(ϕ) indicates the reflectance of the stimulus at an angle ϕ from the centre (in the fixed coordinate system of the arena) and varied linearly from 0.04 (for the black peak) to 1 (for the white peak). (Since photoreceptor cells respond to light, we replaced *X_ink_* with 1 − *X_ink_* to convert the ink value to the intensity of light reflected by the stimulus).

To account for differences between *D. africanum* and *P. lividus*, two parameter values of the model were modified, specifically: the steepness of RN neurons’ sigmoidal response function was set to 2.23 (vs. 3 in the *Diadema* model) while the threshold for the population vector length was set to 1.5 (vs. about 4 in *Diadema* in the presence of a 1st Hermitian stimulus; see Li et al., 2023). All other parameters were as reported in Li et al. (2023). Fixing the positions of the PRCs generated one model of one urchin. Since the results of the model (mimicking the ability of the sea urchin to see the stimulus) depend on the initial orientation of the sea urchin with respect to the stimulus, each model urchin was tested in 200 trials (compared with 70 trials in the experiments). In each trial, the animal started from the centre of the area with a different random orientation (see Li et al., 2023 for details). The ability of the model to detect a stimulus was reported as the success rate, where a trial was deemed successful if the final position of the animal, as inferred from the population vector, was within 72 deg of the centre of the stimulus (36 deg each side; delineated by the green arms in Fig. 3).

**Figure 3.**
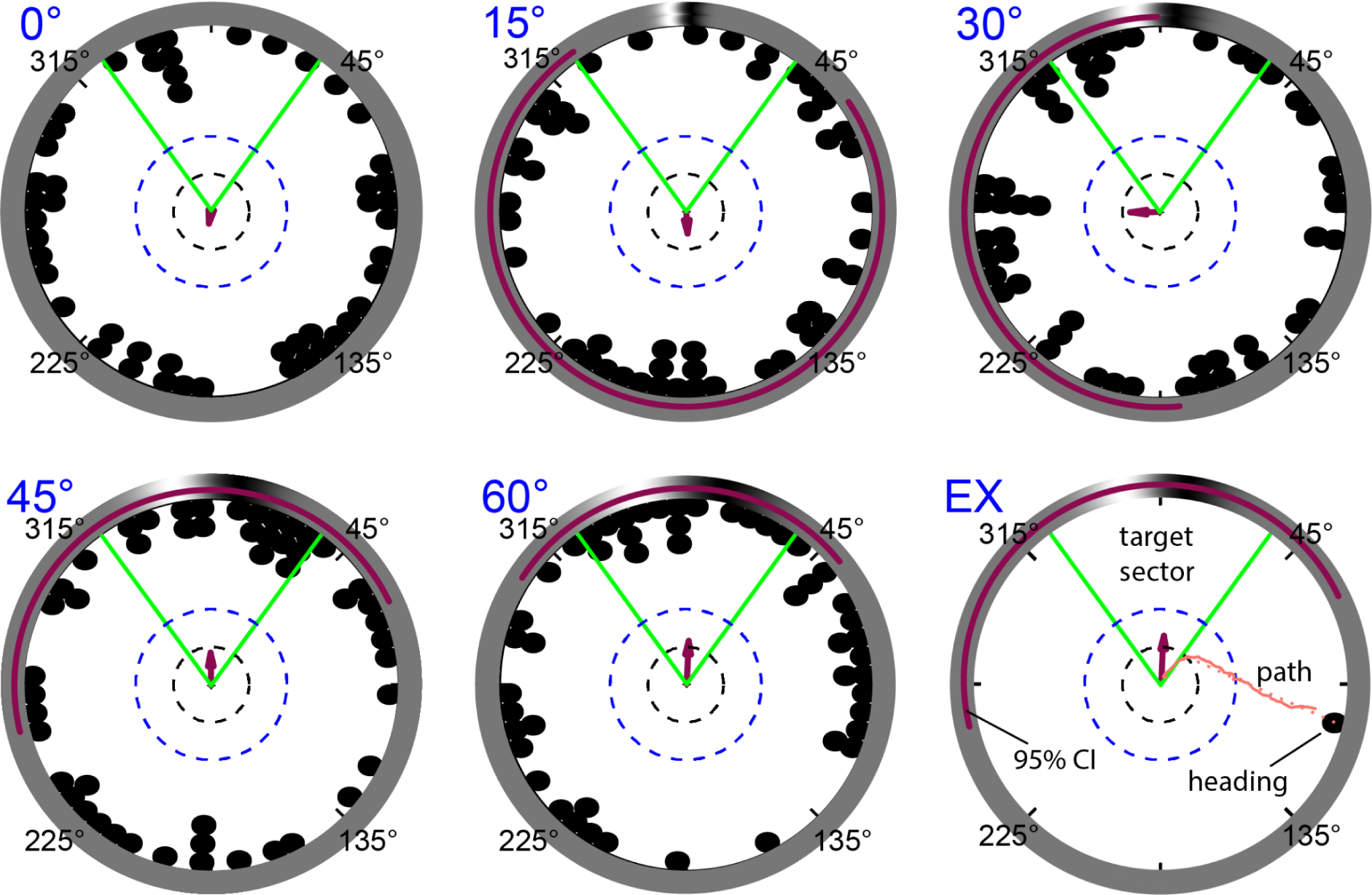
Tracks of *Paracentrotus lividus* in a behavioural test of spatial vision with 1st Hermitian stimuli of 6 differing arc widths, using 50 individuals. All individuals were tested for arc widths of 30 deg to 60 deg whereas 49 were tested for arc widths 0 deg and 15 deg. We calculated headings from the vector made by the animal crossing the dashed circles indicated. Arrows represent the mean resultant length of headings. The green V demarcates the target sector - trials wherein the animal headed towards this sector were classed as successful orientation and vice versa. The red band is a 95% confidence interval (using maximum likelihood estimation) for each condition. The outer circle represents the outer wall of the arena, including the stimulus. The bottom right panel (EX) illustrates a single tracked path and the resultant heading, which was not oriented as it is outside the target sector.

## Results

### Motivation, visual orientation and taxis

We performed taxis experiments in the sea urchin *P lividus* as described in Methods. During initial trials, we used a wide, salient stimulus, of 150 deg, to check for evidence of a response before testing with a range of narrower stimuli. The 150 deg stimulus evidenced only a small effect, when animals were placed on the arena floor centre, as most animals began a straight and apparently arbitrary course to the wall, shortly after placement. However, we observed that on reaching the arena wall, animals frequently settled close to the stimulus, and we sought ways to elicit orientation during trials. Placing the animals atop a clear acrylic cylinder, which partially obstructed their locomotion in all directions, but not their field of view, we found them more likely to orient (Fig. S3), moving slower from the centre, often extending their tube feet for a period, and being more likely to approach the stimulus.

When exposed to the experimental conditions - i.e. moved into the chamber under brighter light, the animals frequently enact what might be an orientation behaviour. They remain in place for upwards of 30 seconds and extend their tube feet fully, directly outwards from each ambulacrum, with buccal tube feet meeting the floor and the remaining tube feet undulating in the surrounding water. They form a wedge with their ambulacral spines leaving a V-shape over the entire ambulacrum into which the tube-feet extend, giving the animal a five or ten-pointed ‘stellate’ appearance. Each ambulacrum thus has a distinct ‘field of view’ around the horizon, delimited by a dense row of ambulacral spines.

Occasionally, animals (especially the largest individuals) will linger at the arena centre for the trial duration, but generally, they crawl at a steady pace on a relatively straight course. While moving, they frequently maintain these V-shapes around the one or two ambulacra nearest the leading edge of the animal, continuing to extend their tube feet outwards from these. On reaching the arena wall, the extended tube feet make contact and the animal draws its body to the wall, sometimes climbing upwards and settling on it.

The extended tube feet undoubtedly have non-visual sensory roles, including those related to identifying nearby objects. But the V shape of spines around the ambulacrum might screen off-axis light to facilitate vision. This posture could therefore function to augment vision, and improve resolution.

All animals appeared to be in good physical condition throughout the experiments: Without the spine loss or torpid posture of ailing sea urchins.

### Resolution

A total of 248 trials were successfully tracked across five increasing stimulus widths, including a negative control with only a grey background (0 deg), with at least 49 individuals at each level (Fig. 4 & Table 1) from 50 individuals. Thus, the trials per stimulus level are balanced and independent. For the three narrower stimuli (0 - 30 deg) no orientation was evident, whereas for the wider stimuli, we find evidence of orientation (see table 1).

**Figure 4.**
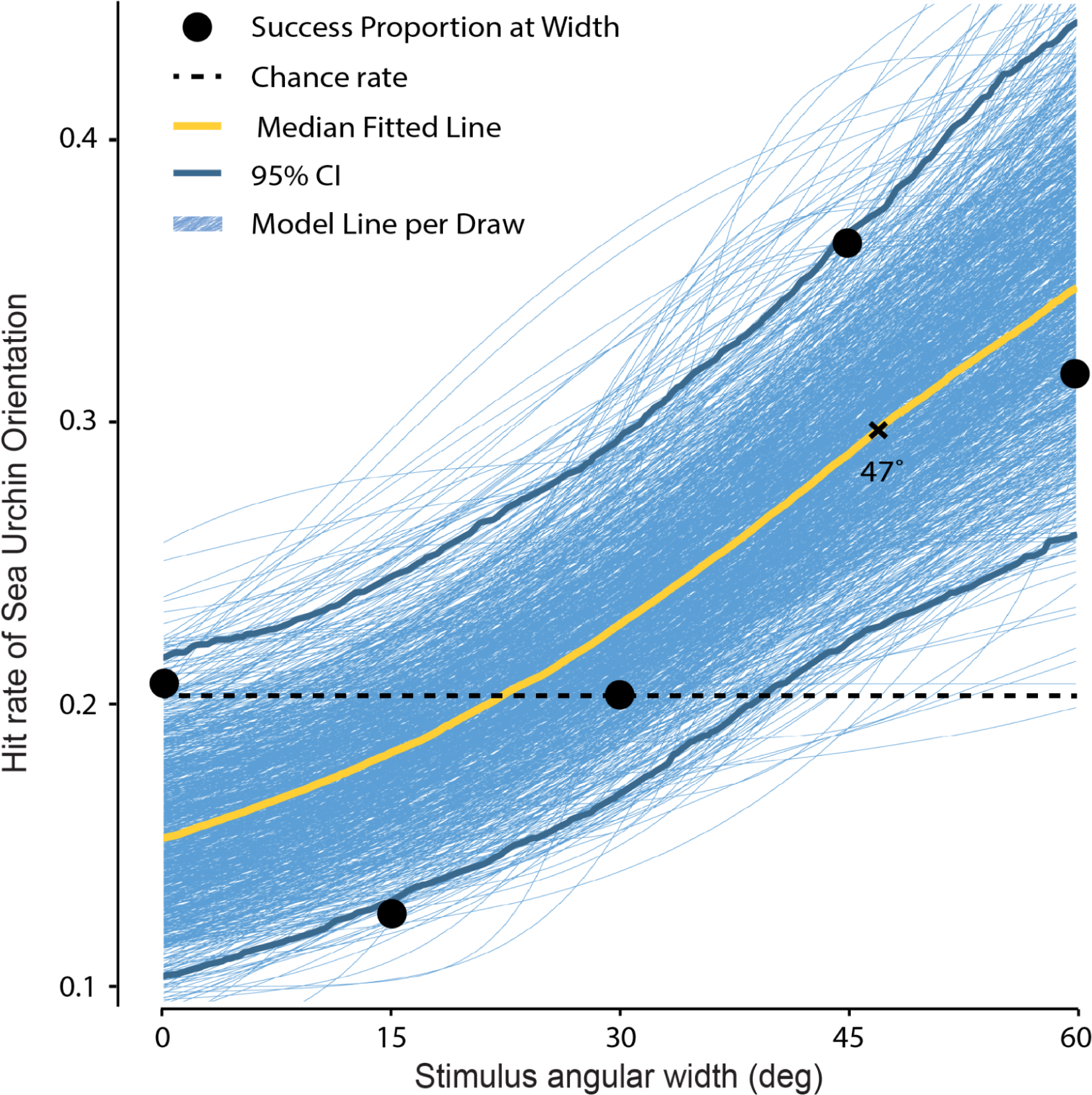
Bayesian model of the psychometric function used to find the spatial resolution exhibited by *Paracentotus lividus*. The plot shows the proportion of success of sea urchin orientation (hit rate) against the angular width of the visual stimulus estimated by the model. All 50 individuals were tested for arc widths of 30 deg to 60 deg whereas 49 were tested for arc widths 0 deg and 15 deg. The black dots represent the proportion of successes at each tested arc width. The blue sigmoids represent the model formula at each sampled iteration over the range of angular widths of the visual stimulus; the yellow line represents the median. The psychometric function is bounded between two asymptotes, which are also estimated: a lower asymptote indicating the random chance rate and an upper asymptote indicating the lapse rate (failure to orient even with a salient stimulus). Model line per draw represents the fitted line drawn from the posterior values for 4 000 iterations across the four Markov chains. The black dashed line (chance rate) is the proportion of successes that would be expected by random chance (0.2).

**Figure 5.**
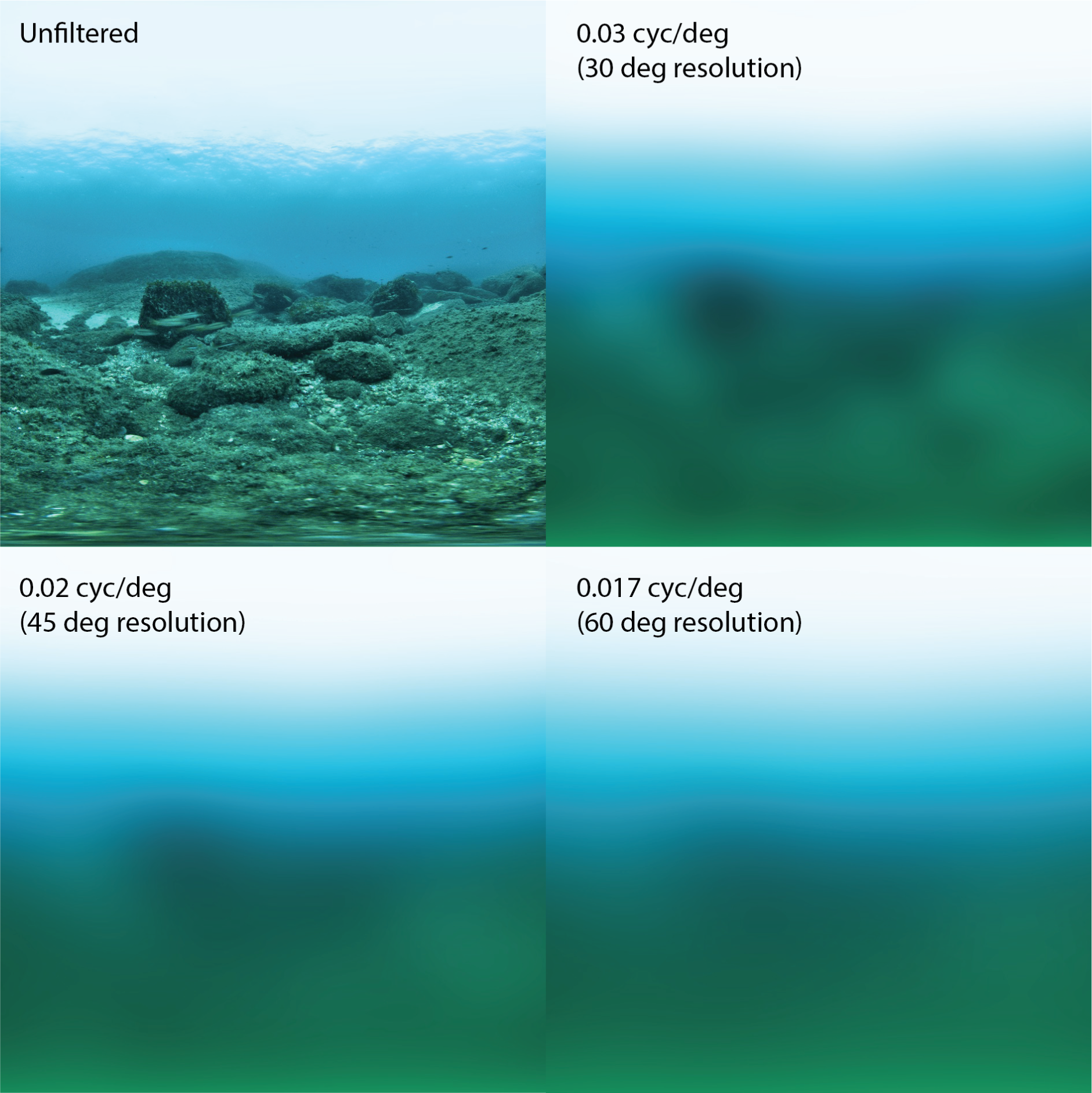
Photograph of *Paracentrotus lividus* habitat at 10 m depth in the bay of Naples (top left). The scene is taken with a 180 deg fisheye lens, and the resulting circular image has been remapped to linear vertical and horizontal coordinates. The remaining panels show this image filtered via Gaussian blurring to represent the information loss by a visual system at three progressively worse resolutions. The median estimate of resolution for *P. lividus* (47 deg) is close to that of the 45 deg panel. Note that the colour images only illustrate visual acuities in the range of sea urchin vision: sea urchins do not see the colours.

**Table 1:** Summary of orientation response by *Paracentrotus lividus* in behavioural trials across five stimulus arc widths. Mean heading is the circular mean of the headings per condition. A binomial test tests the null hypothesis that the proportion of animals orienting towards the stimulus is equal or less than random. FDR-adjusted p-value is that of the binomial test adjusted by the false-discovery rate (Benjamini & Hochberg, 1995).

| Arc width (deg) | Mean Heading (deg) | Mean Resultant Length ($\rho$) | Successes (n) | Total (n) | Binomial p-value | FDR adjusted p-value |
| --- | --- | --- | --- | --- | --- | --- |
| 0 | 190 | 0.07 | 10 | 49 | 0.528 | 0.695 |
| 15 | 177 | 0.12 | 6 | 49 | 0.945 | 0.945 |
| 30 | 270 | 0.16 | 10 | 50 | 0.556 | 0.695 |
| 45 | 359 | 0.18 | 18 | 50 | 0.006* | 0.03* |
| 60 | 3 | 0.26 | 16 | 50 | 0.031* | 0.078 |

The mean resultant length (ρ) is a measure of orientedness, ranging from 0 to 1 for complete directedness (i.e. if all paths had identical headings). The one-tailed binomial tests found significant evidence for orientation only for the two widest stimuli (45 and 60 deg). To avoid multiple-testing, a false discovery rate correction was applied (Benjamini & Hochberg, 1995), upon which only the 45 deg remained significant.

To ensure that the animals were orienting and find the resolution, we relied primarily on a Bayesian probabilistic model. The Bayesian probabilistic model converged with effective sample size >10% of sampled draws and Gelman-Rubin statistic (Ȓ) below 1.1 (Gelman & Rubin, 1992).

Fig. 4 indicates 1000 sampled draws from the Bayesian regression model in blue. The median estimate is shown in yellow. The liminal threshold is derived from the highest rate of change: the inflection point of the sigmoid. The sigmoid is bounded between two asymptotes, which are estimated in the statistical model. We carried out a prior predictive test to test to what extent the priors impact the result. This test (Fig. S4) supports the model as it shows that the priors did not guide the model towards evidencing spatial resolution and were somewhat conservative. A leave-one out model comparison tested whether the model explained the variation better than a null model which lacked the stimulus width predictor, and this was the case (ELPD diff -2.3, SE diff 2.7).

To illustrate the visual acuity estimated for *P. lividus* in these experiments, we downsampled the resolution of an image of their habitat via Gaussian blurring to demonstrate the visual information available to *P. lividus* (Fig. 6).

*P. lividus* vision almost certainly lacks colour discrimination, having only a single known R-opsin (D’Aniello et al., 2015). Moreover, the resolution we have estimated is for across the horizon, given our behavioural paradigm, and omits the vertical oral-aboral plane. In a well lit environment, the best estimate of resolution for *P. lividus* suffices to detect a cluster of boulders (a structured environment, potentially representing shelter) against the open ocean floor at several metres distance.

Additionally, we downsampled the resolution of images of one *P. lividus* individual using AcuityView (Caves & Johnsen, 2018) to simulate this visual acuity at increasing distance (Fig. S6). This resolution suffices to detect conspecifics from the same distance against a bright background.

### Neurosensory computational model

To understand the neural basis of the observed orienting behaviour in *P. lividus*, we tested a previously published model of visual orientation in sea urchins (Li et al., 2023) against the data collected here. This model uses known anatomical and physiological components of the nervous system of sea urchins (Fig. S5). It was originally proposed to understand visual orientation in the sea urchin *Diadema africanum* by explaining how decentralised vision can emerge from the neural integration of the light information coming from the surface of the animal’s body (Kirwan et al., 2018; Li et al., 2023). The model’s PRCs are distributed on the tube feet along the ambulacra and relay visual information to neurons in the RNs of the same ambulacra. The RN neurons project to neurons in the ONR, which integrates the neural signals coming from several ambulacra to produce detection of visual stimuli and movement towards (or away from) the stimuli (see Methods and Fig. S5).

We show in Fig. 6 that the model successfully recapitulated the visually-driven behaviour of *P. lividus* in the presence of the same stimuli as those used in the behavioural experiments (compare Figures 3 and 4 with Fig. 6). Starting from the *Diadema* model, it was sufficient to modify only 2 parameters to capture the *P. lividus* data, suggesting that the basic anatomical and physiological structure of the model may generalise to decentralised vision in a wider range of echinoderm species.

**Figure 6.**
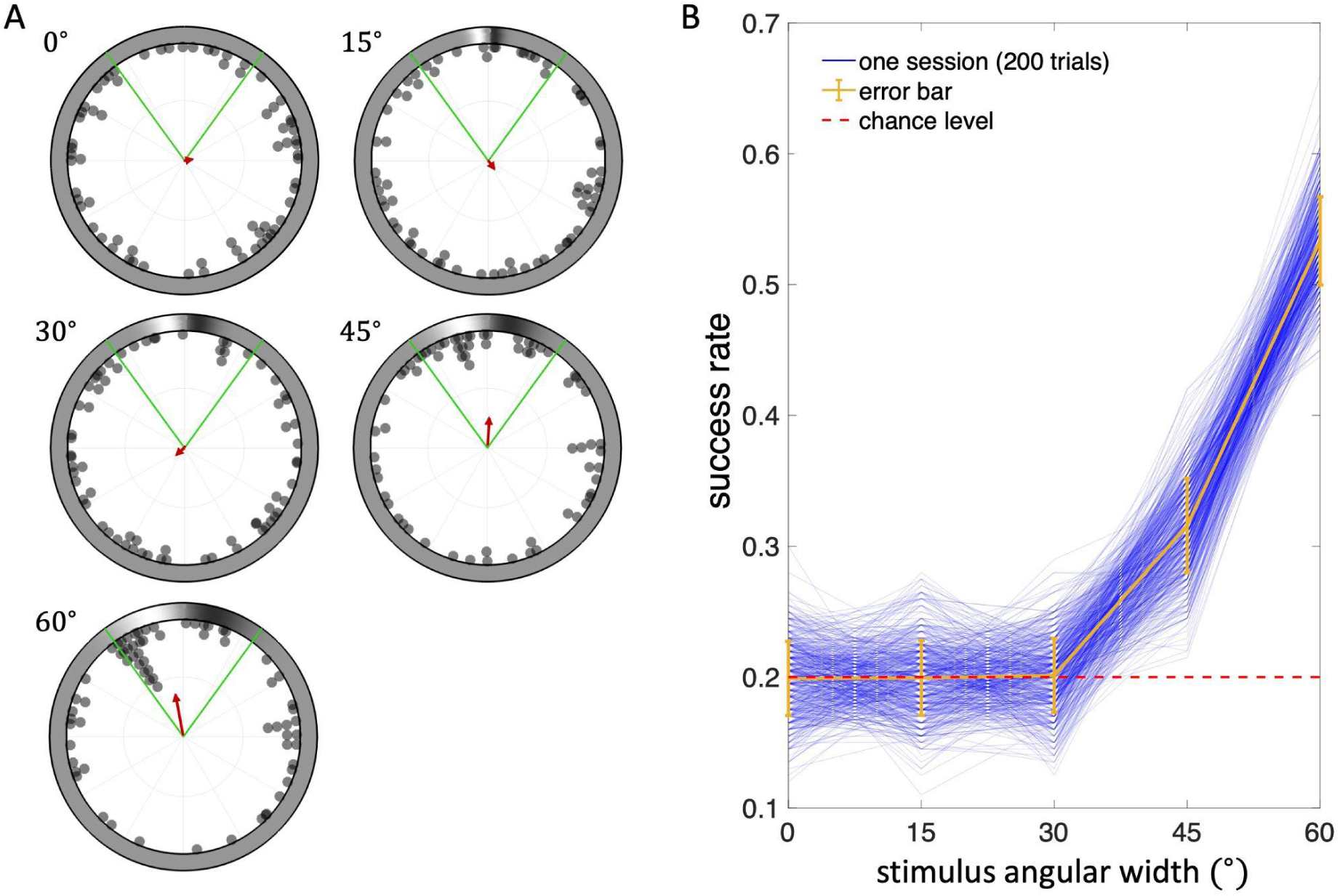
Results from the neurosensory computational model. A. Tracks of *P. lividus* model in simulations of behavioural test of spatial vision with 1st Hermitian stimuli of 5 differing widths (compare with Fig. 3). Fifty model animals (two trials per animal) were used in these trials. Headings were calculated based on the population vector (see the Methods). B. Success rate vs. angular width for 200 sessions (=model animals; compare with Fig. 4). A trial was successful if the final position of the animal, as inferred from the direction and strength of the population vector, fell within the green arms of panel A (see Methods). Each blue line represents a session with one model animal (200 trials per session). Yellow: average success rate across sessions. Error bar: standard deviation of success rate across sessions.

## Discussion

### Low resolution vision may aid *P. lividus* to find structured habitats and shelter

Wild *P. lividus* inhabits rocky areas and seagrass beds (such as *Posidonia*) and avoids exposed areas (Boudouresque and Verlaque, 2013), such as sandy bottoms, which often divide these more structured habitats. Like many sea urchins, *P. lividus* adults in the Mediterranean typically forage at night and shelter during the day. Populations inhabiting areas of low predation, in contrast, show a reduced sheltering behaviour (Sala & Zabala, 1996). Individuals found in open areas lacking rock boulders cling to stones, shells or other scattered objects.

Finding shelter and remaining within structured habitats, or returning to it if displaced nearby, likely constitutes an important visual task for sighted sea urchins (Kirwan et al. 2018) and is the behavioural paradigm which our experiments aimed to recapitulate. One role for the visual taxis response we elicited could be to find shelter when displaced by strong currents. Aside from camouflage, rocks provide solid purchase to which sea urchins can cling, especially when wedged within a cleft or crevice. This conceals their vulnerable underside and stymies predators attempting to dislodge them.

In our trials, the largest individuals occasionally remained near the arena centre, suggesting that they are less motivated to shelter when moved and exposed to bright light (which was also the case for the largest *D. africanum* (Kirwan et al. 2018)). The major predator of *P. lividus* in the Mediterranean, the seabream *Diplodus sargus*, does not typically feed on *P. lividus* of this larger size class, with a test wider than 50 mm (Sala, 1997; Guidetti, 2004). A comparable observation has been made in arena trials of large *Diadema* individuals, which were less likely to approach visual targets than their smaller conspecifics (Kirwan et al. 2018). Both findings point towards a considerable role of internal motivation in sea urchin visual responses. *P. lividus* needing a physical obstruction to exhibit significant object taxis adds to those considerations.

Visually locating rocky reef habitat structures (Fig. 5) may also help localise food resources. Under low predation pressure, sea urchin populations can explode (Sala & Zabala, 1996), leading to overgrazing and a depletion of macroalgae and seagrass in their habitat (Eklöf et al. 2008). Sea urchins can feed on biofilms (Porter et al. 2019, Kasper 1992) and, absent predators, often aggregate on rocks in large numbers (Sala & Zabala, 1996).

Two species of seastar (which are echinoderms) locate food resources through vision, using simple discrete eyes at the end of each arm, to prey upon corals. They have somewhat sharper vision than *P. lividus*, estimated at 15-30 deg for *Linkea laevigata* (Garm and Nilsson, 2014) and 8 deg for *Acanthaster planci* (Petie et al., 2016). The low resolution vision of *P. lividus*, although much coarser than that of its seastar cousins, might also help orient towards food-rich areas.

According to the prediction of the visual filtering (Fig. S6), *P. lividus* may recognize another sea urchin from a distance of 45 cm or more, where contrast and illumination suffice (e.g., against a light-coloured surface in daylight). The coarse resolving power predicted by our experiments and the external sexual monomorphism in sea urchins precludes a visual component to sex recognition.

Based on their size, the animals used in these trials were likely older than three years (Boudouresque and Verlaque). In the lab, *S. purpuratus* exhibits simple negative phototaxis approximately one month after metamorphosis (Ullrich-Lüter et al. 2011), when the calcite skeleton forms, rendering the body opaque. The tube feet of *P. lividus* express R-opsin as early as the late larval rudiment, before metamorphosis (Paganos et al. 2022). Yet, it is unclear at what size juvenile sea urchins are capable of vision.

### Using isoluminant stimuli to test for resolving vision

By decoding an isoluminant biphasic signal in the form of the visual stimulus, *P. lividus* has demonstrated image-forming vision in these experiments, albeit of meagre resolving power. Vision has been reported in several species of Echinacea (the suborder which includes most ‘regular’ radially-symmetrical sea urchins, and excludes diadematoids) based on behavioural experiments.

Some of these avail of bar or spot stimuli, which could be detected with an acuity worse than proposed due to the locally greater intensity difference. Such a stimulus could theoretically be detected by a single receptor with a wide acceptance angle, being swept over it and registering changing intensity over time. Spatial (image-forming) vision involves simultaneous sampling from multiple sets of directions to compare intensities (Nilsson, 2009). Our reliance on an object taxis task (which requires spatial vision) thus confirms image-forming vision in this group.

Kirwan et al. (2018) sought narrowband signals applicable to directional tasks, such as wavelets with localised optima to orient towards. Continuous wavelets performed best, especially a difference of Gaussians. As Kirwan et al. (2018) found that isoluminant stimuli comprising continuous signals are advantageous, we did not seek to use a discrete bar in this case. We instead sought a signal with comparable properties to the difference of Gaussians (DoG) previously applied, but which is a dual wavelet, balanced about the zero-crossing with a peak and trough of equal amplitude.

### Probabilistic modelling facilitates behavioural studies where animals lapse

The Bayesian modelling of the psychometric function is a powerful approach, which guards against separation in logistic regression (Gelman, Jakulin, Pittau, & Su, 2008), allows for many parameters and nonlinear models to be estimated effectively, and returns rich probabilistic information. Kirwan et al. (2018) with regard to vision in *Diadema* applied a logistic regression model approach to measure the acuity of the spine-pointing response but did not to estimate spatial vision based on a locomotion task similar to these experiments, as there were too few stimulus levels.

Including several stimulus levels allows for the psychometric function to be implemented, which can make efficient use of the data and avoid multiple-testing. Bayesian estimation allows us to identify even small effects and describe them as probabilities. The nonlinear approach, estimating chance rate and lapse rate asymptotes outside the psychometric function, allows us to estimate the lapse rate in particular (the rate at which the animal does not respond to even a salient stimulus), which is high in this case. By ensuring that priors do not bias the model towards a given conclusion, and by checking the model validity, we can investigate behaviours even in cases where a robust response is lacking.

### A model of decentralised vision, developed for Diadematoida, accords with *P. lividus*

The orientation behaviour of *P. lividus* via object taxis is quantitatively explained by our recently published model of decentralised vision in echinoderms (Li et al., 2023). The model was proposed to account for the visual orienting behaviour of *D. africanum* reported in Kirwan et al. (2018). It provides a mechanistic explanation by combining a particular process of visual integration and readout at the level of ONR neurons with general anatomical characteristics of the sea urchin nervous system. Detailed information about the model can be found in Li et al. (2023). Here, we briefly summarise its salient features and discuss their viability in light of the available experimental data.

The model assumes that light information is captured by PRCs distributed over the animals’ epidermis around the ambulacra, and that PRCs relevant for object resolving vision are rhabdomeric opsin expressing PRCs located in skeletal depressions at the base of each tube foot (Ullrich-Lüter et al. 2011). Whereas R-opsin+ PRCs in *Diadema* have so far not been detectable on RNA or protein level (Sumner-Ronney & Ullrich-Lüter 2022), R-opsin+ PRCs in *P. lividus* have recently been shown to express various retinal marker genes in early developmental stages of this sea urchin (Paganos et al. 2022), making them suitable candidates for the task of resolving vision.

Starting from PRCs distributed over the animal’s body, the model accounts for three main layers representing the flow of information from PRCs to RN neurons to ONR neurons (Fig. S5). It includes recurrent connections inside ONR neurons and among neurons in the same RN, in addition to feedforward connections between the layers (PRCs to RN neurons and RN neurons to ONR neurons). When PRCs on the animal’s tube feet are stimulated by light, RN neurons in the same ambulacrum are inhibited. RN neurons then indirectly suppress excitatory neurons in the ONR (eONR) so that, when PRCs are activated by light, the eONR neurons are overall excited by the stimulus. The eONR’s lateral recurrent connections enable the integration of data from adjacent ambulacra. The eONR’s overall activity is readout as a “population vector”, whose length determines whether the stimulus has been detected, and whose direction determines the location of the centre of the stimulus (see Methods).

The viability of each of these proposed pathways and mechanisms in light of the available experimental data was examined by Li et al. (2023). The existence of PRCs on the tube feet of sea urchins has been established in various sea urchin species (reviewed in Sumner-Rooney & Ullrich-Lüter 2022), and there is considerable clear evidence for a pathway from RN neurons to ONR, in addition with evidence for the integrative role of the ONR for both stimulus integration and coherent movement (Yoshida, 1966).

We chose a double-inhibitory pathway from PRCs to RN neurons and from the input state of ONR neurons to eONR neurons to account for lesion experiments in other species of sea urchins reported by Yoshida et al. (1984). Thus, we tried to maximise the available data that could be captured by our model, especially in view of the paucity of experimental data. In particular, it is not clear whether PRCs, or which neurons in echinoderms, mediate excitatory or inhibitory action on their projections.

The fine structure of neural organisation and morphology is largely unexplored in sea urchins, and despite various neurotransmitters and synaptic proteins being expressed in neuronal tissues of developmental stages of *P. lividus* (Paganos et al. 2022) it is unclear if and how neurons in the sea urchin communicate via chemical synapses (Florey & Cahill, 1980; Kawaguti, 1964). Absent more detailed experimental evidence, we have combined known anatomical features of sea urchins with plausible hypotheses for their functional organisation. The result is a mechanistic model that allows us to make predictions on which visual stimuli can be resolved using a decentralised sensory-visual-motor architecture in these animals (Li et al., 2023).

The model only required minimal modifications to account for the orienting results obtained here in *P. lividus* – specifically, it required changes to two parameter values. The first one, the threshold length for the population vector, determines whether or not an extended stimulus has been detected. We had to reduce this parameter value from about 4 (in *Diadema* in the presence of 1st Hermitian stimulus) to 1.5 (in *P. lividus*), which means that a lower threshold for visual detection is required for *P. lividus*. Yet, *P. lividus* need not have greater visual acuity. Assuming that both species have a nervous system well characterised by the model, *P. lividus* may require lesser neural activity to resolve a given stimulus. The second parameter modification was a slight reduction of the steepness of the sigmoidal response function of the RN neurons (from 3 to 2.23). The sigmoidal response function gives the output activity of RN neurons as a function of the PRC input; a reduction in steepness gives a slightly more nuanced response to the same input.

Given the differences in morphology between *D. africanum* and *P. lividus,* it is unsurprising that parameters such as signal detection levels may vary according to different ecological needs of the animals. While the tests of adult animals are in a similar size range, *D. africanum* has a thin skeleton but long and hollow spines, and the longest primary spines are particularly sharp, brittle and poisonous. *P. lividus* features a more robust skeleton, but much shorter, solid spines.

Defence behaviours also differ between these species. Under predation, both species typically lodge themselves within crevices or in shelter during the day, often aggregating to present a formidable phalanx of spines to a prospective predator. This could be especially valuable for juveniles, which shelter among larger adults (Ouréns et al., 2014). Both species tend to congregate in shaded parts of a tank. However, *Diadema* has an active, photoreceptive response to a potential predator: Waving its long, poisonous spines towards a looming shadow. *P. lividus*, like most sea urchins, lacks this startle response. *P. lividus*, in some parts of its range, also burrows an indentation into the rock surface to help lodge itself.

The core model is able to correctly predict behavioural outcomes in both species, *D. africanum* and *P. lividus*, despite their distinct morphologies and photic behaviour, which supports the robustness of its underlying neurobiological assumptions. The model, being able to successfully bridge species-specific differences, even for phylogenetically distant species, allows us to draw more generalised conclusions on sea urchin resolving vision.

### Limitations of the study

The visual acuity of *P. lividus* could be better than these behavioural experiments showed, if the animals were unmotivated to approach visible targets. A detectable target may not be relevant for them or amenable to testing with the behavioural paradigm used. While animals appeared in good condition and had been accommodated to the housing conditions before the experimental trials, the housing or experimental conditions (e.g. lighting regime) may not have been optimal for the experimental paradigm and impacted their visual capability or mood, so as to make them less inclined to orient.

Behavioural experiments with a greater array of stimulus types would enrich the neurosensory model of decentralised vision, by verifying its predictions and allowing for improvements. This could include both discrete and continuous visual stimuli among several species.

### Questions for future experiments

Various sea urchin species have demonstrated orientation towards visual targets, and despite potential difficulties given the target test designs, might thus possess coarse vision. However, of those tested, only two, *S. purpuratus* (Sodergren et al. 2006) and *P. lividus* (Marlétaz et al. 2023), are available for functional genetic experimentation, which is essential to obtain final proof of the mechanisms involved in sea urchin vision. Further experiments behaviourally testing the acuity of the *P. lividus* visual system and refining the theoretical model will provide a framework for integrating and testing future findings on its structural and molecular composition.

### Conclusions

Regular sea urchins can visually resolve objects and orient towards them. Their taxis reactions towards different target types might depend on the angular sensitivity of their species-specific photoreceptor system, as well as on motivational states influenced by physiological factors such as individual body size in relation to their predators’ prey pattern.

## List of Symbols and Abbreviations

DoG: Difference of Gaussians
eONR: Excitatory ONR neurons
ONR: Oral nerve ring
PRC: Photoreceptor cell
RN: Ring nerve

## Acknowledgements

We thank Michael J. Bok for assisting with collecting ELF images and processing them. We thank Davide Caramiello for his assistance with animal husbandry and Francesco Izzo for photography.

## Author Contributions

The project was conceived by MIA, DEN, JUL and GLC. The behavioural experiments were designed by JDK in conjunction with all of the authors. Animal collection and husbandry facilities were provided by MIA. JDK built the setup and carried out the behavioural experiments and statistical analysis. Neurosensory modelling was carried out by TL and GLC. Writing and editing was led by JUL, GLC, and JDK and all authors contributed.

## Data availability

Custom MATLAB code to simulate the neurosensory computational model is freely available at https://github.com/lacameralab/diadema. Custom MATLAB code to track animals in recordings is available at https://bitbucket.org/jochensmolka/dtrack.

